# Sexual selection driven by direct benefits leads to the erosion of direct benefits

**DOI:** 10.1101/2025.11.07.687154

**Authors:** Jana M. Riederer, Giulia Cordeschi, Marta Mosna, Franz J. Weissing

## Abstract

Most sexual selection models assume that the evolution of female choosiness is driven by indirect (genetic) benefits, such as the production of more viable offspring or more attractive sons. There is ample empirical evidence that female choosiness can also provide direct (non-genetic) benefits, including access to good territories or paternal care. Yet, theoretical models of direct-benefits sexual selection are scarce. Here, we use individual-based evolutionary simulations to explore the joint evolution of female mating preferences and male ornamentation under sexual selection driven by direct benefits. We find that, within each generation, male ornamentation reliably indicates paternal quality (the direct benefit); females thus benefit from being choosy and readily evolve preferences for male ornamentation. Yet this in turn selects for males to invest resources in ornamentation at the expense of paternal care. Across generations, female selection for ornamented males (driven by the pursuit of direct benefits) thus erodes the very benefit it seeks, leading to population decline and potentially extinction. Our results highlight the complexity and non-intuitive nature of direct-benefits sexual selection.

## Introduction

Sexual selection has fascinated biologists ever since Darwin (1871)^1^ for its ability to generate impressive morphological and behavioural diversity. It can be a driving force in evolution^2^, shape biodiversity^3^, accelerate adaptation^4,5^, and facilitate speciation^6^, but also act as a “double-edged sword” that constrains evolution^7,8^ and drives extinction^9^. It often involves complex coevolutionary dynamics between males and females, fuelling rapid evolutionary change^10^. However, the mechanisms underlying these complex dynamics are not always well-understood.

Generally, sexual selection models consider two types of benefits that choosy females (or males, in sex-role reversed systems) can obtain from non-random mating. On the one hand, by being choosy, females may increase their lifetime reproductive success by increasing their own viability or fecundity. In this case, choosy females gain “direct benefits”, e.g. due to courtship feeding or nuptial gifts, access to a higher-quality breeding territory, anti-predator behaviour of the mate, access to paternal care, or simply the absence of directly transmitted diseases^11–15^. On the other hand, by being choosy, females may not produce more but “fitter” offspring, that is, offspring with a higher reproductive value. In this scenario, choosy females gain “indirect benefits”, e.g. due to “good genes” enhancing offspring viability contributed by their mate, or due to attractive fathers producing fecund “sexy sons”^16–18^ (reviewed in^19,20^).

A large body of theoretical work has aimed to better understand the process of sexual selection^20,21^, but, notably, most of this work has focused on indirect benefits. This may be because sexual selection driven by indirect benefits is considered to be more challenging to explain^22–24^. For example, genetic benefits are not directly observable, raising the question of what ensures signal honesty, and runaway processes driven by indirect benefits may decrease population viability. In contrast, sexual selection driven by direct benefits appears conceptually more straightforward, as benefits such as nuptial gifts or the quality of a male’s territory can often be assessed with relatively high reliability, and the costs and benefits of female choosiness are both expressed in terms of the female’s own lifetime reproductive success, allowing them to be compared more easily. Consequently, one would predict that under direct-benefits sexual selection, female preferences should simply evolve in a way that maximises female fecundity or, more broadly, lifetime reproductive output^25–29^. Given this apparent conceptual simplicity, it is perhaps not surprising that theoretical models addressing sexual selection driven by direct benefits are by far outnumbered by those focusing on indirect benefits, even though, in natural systems, direct benefits may play a more prominent role than indirect benefits in shaping female mate choice^13,30–32^.

However, direct-benefits sexual selection may be less straightforward than it appears^19,20^. First, females may often not be able to reliably assess the quality or magnitude of the direct benefits they are likely to receive. For example, at the time of mating, a female may be unable to predict the level or quality of paternal care a male will provide later on. Therefore, signalling also plays a critical role in the context of direct benefits, raising a parallel question to that posed in the context of indirect benefits: how to ensure signal honesty? A further complication in the context of direct-benefits sexual selection is that male signals may actively compromise the males’ ability to provide direct benefits, such as paternal care. For example, the red nuptial colouration of male three-spined sticklebacks may signal the male’s condition and, hence, the male’s intrinsic paternal quality, but it may also attract predators to the male’s nest. The idea that signalling parental care may compromise parental care has been previously discussed in the literature^33^, but, to our knowledge, has never been investigated systematically.

Here, we present a direct-benefits sexual selection model where males differ in their ability to provide direct benefits (e.g., paternal care), and females can evolve preferences for male ornamentation that might indicate the benefits to be expected. In response to these preferences, males can evolve ornaments. However, ornaments are costly in that they trade off against paternal care (i.e. against the direct benefit provided): in other words, ornaments carry a fecundity cost. Note that we are here using parental care as an illustrative example of a possible direct benefit of mate choice, but the model can also be applied to other forms of direct benefit. To our knowledge, all current models of direct-benefits sexual selection are phrased in terms of quantitative genetic^25,34–36^ or population genetic^25,26,29^ equations. In contrast, we here use individual-based evolutionary simulations. While all approaches have their pros and cons (see ^20^), we chose a simulation approach because it incorporates key components of evolutionary processes (such as individual variation, polymorphisms, and stochasticity) more easily than other approaches and allows us to study the evolutionary dynamics close to and away from equilibrium. Previous studies have demonstrated that such features of individual-based simulations lead to novel insights into the complex dynamics of co-evolving traits in males and females (see, for example,^37–40^).

With this model, we address the following questions: Under what conditions do females evolve preferences for male ornamentation that signals direct benefits? When do males evolve ornaments that are related to the amount of direct benefit (e.g. paternal care) they will provide? Are these signals honest? Do the evolutionary dynamics lead to equilibrium, and what is the evolutionary outcome regarding the trade-off between paternal care and ornamentation?

Our findings will also shed new light on the long-standing question of whether and when one should expect that male ornamentation is positively or negatively related to paternal care. For example, Hoelzer’s (1989)^41^ “good parent process” is based on the idea that sexual ornamentation is a reliable signal of a male’s parenting ability; hence, it predicts a positive correlation between ornamentation and paternal care. Conversely, a negative correlation is predicted by many variants of the “differential allocation hypothesis”, which assumes that individuals adapt their parental effort to their own attractiveness or the attractiveness of their mate^42,43^. Strikingly, in our model, both positive and negative correlations can emerge, highlighting the dynamic and context-dependent nature of the relationship between ornamentation and parental care.

## Results

The structure of our model is illustrated in Figure 1. We consider a population of males and females. Each male has a certain amount of resources available; the amount varies between individuals and is determined stochastically. Each male can invest a proportion *S* of his resources into producing an ornament; the remaining resources are invested in the direct benefit, i.e., in paternal care. Variation in resource level between males depends solely on the environment, whereas variation in ornamentation and paternal care is driven by both the environment and the males’ investment strategy *S*. During the mating season, females encounter males, and can choose to accept or reject potential mates, based on how well the males’ ornament size matches their preference *P*. However, time is limited: females that have not found a sufficiently “attractive” mate within a certain number of tries are “out of time” and mate with a random male. After the mating season, the resulting offspring are reared by the parents, where offspring receiving more paternal care have a higher survival probability. We allow both the male investment strategy *S*, and the female preference *P*, to evolve in two model versions that differ in whether population size is fixed or changing dynamically.

**Figure 1:**
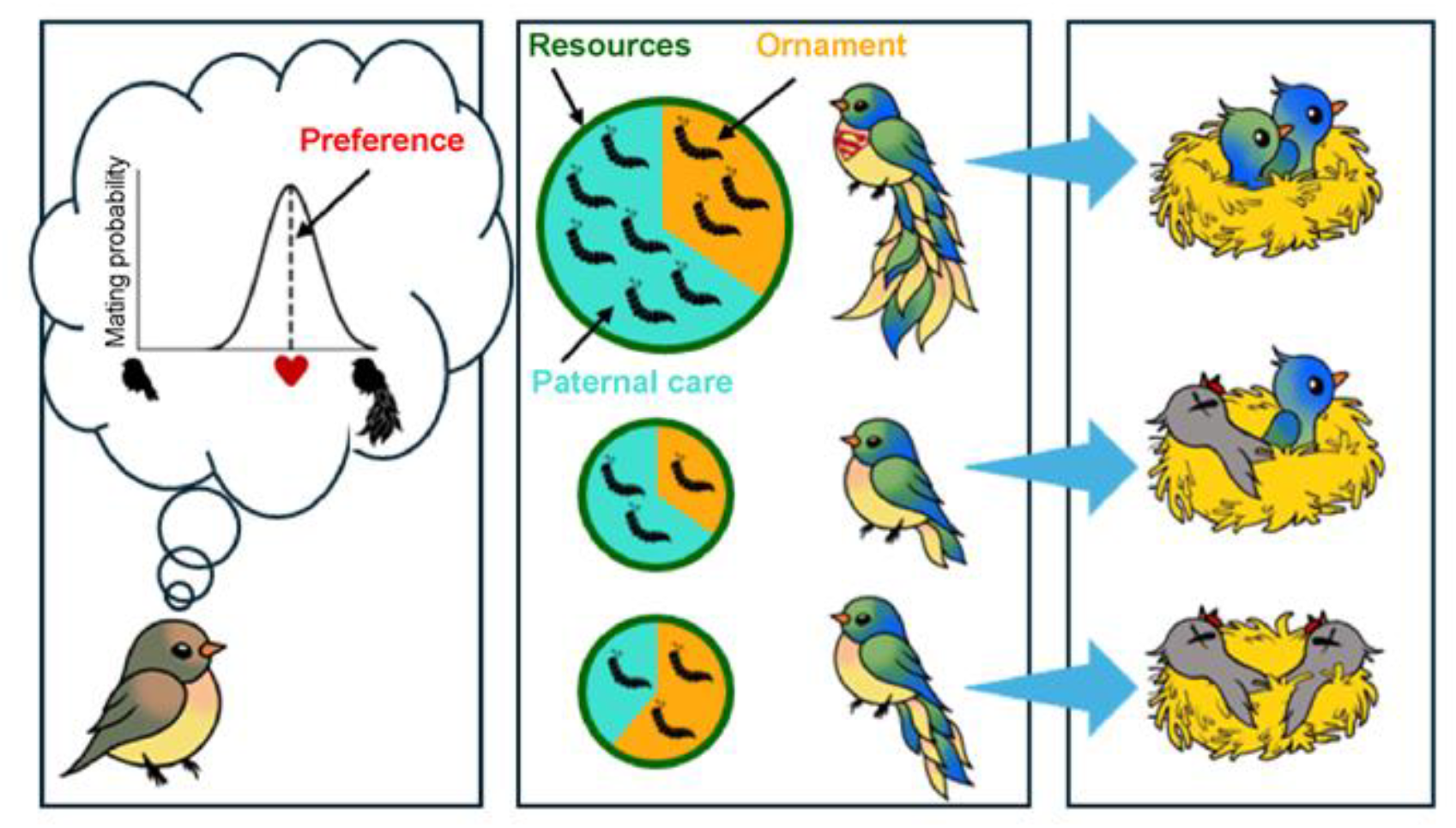
The model in a nutshell. Females are more likely to mate with a male whose ornament matches their preferences. Males differ in how many resources they have acquired (the size of the “pie”); they can invest these partly in ornamentation and partly into providing direct benefits (e.g. paternal care), depending on their investment strategy. Offspring survival depends on the level of direct benefit received. Note that the third male invests a greater proportion of his resources in ornamentation than the first male; however, as he has fewer resources available overall, the resulting ornament is nevertheless smaller.

### Sexual selection erodes direct benefits

We first consider our baseline scenario, in which male resource levels are drawn from the interval [0.5,1.5] and females have time to encounter at most 10 males (i.e. females can reject up to 9 males before mating with the randomly chosen 10^th^ male). Figure 2 shows the outcome of this baseline scenario: evolution of investment in (and preferences for) large ornaments, trading off against investment in paternal care. Starting from a population of unornamented males (i.e. *S* = 0, all resources are invested in paternal care) and females with a preference for unornamented males (i.e. *P* = 0), we find that sexual selection dynamics is readily established: females evolve preferences for larger ornamentation; correspondingly, males evolve to invest in larger ornaments. However, this is accompanied by a decrease in paternal care, i.e. a decrease in the direct benefit that is honestly signalled (see below) by these ornaments: increased ornamentation has evolved at the expense of paternal care, since ornamentation and paternal care trade off against each other. In other words, direct-benefits sexual selection here erodes the direct benefit.

**Figure 2:**
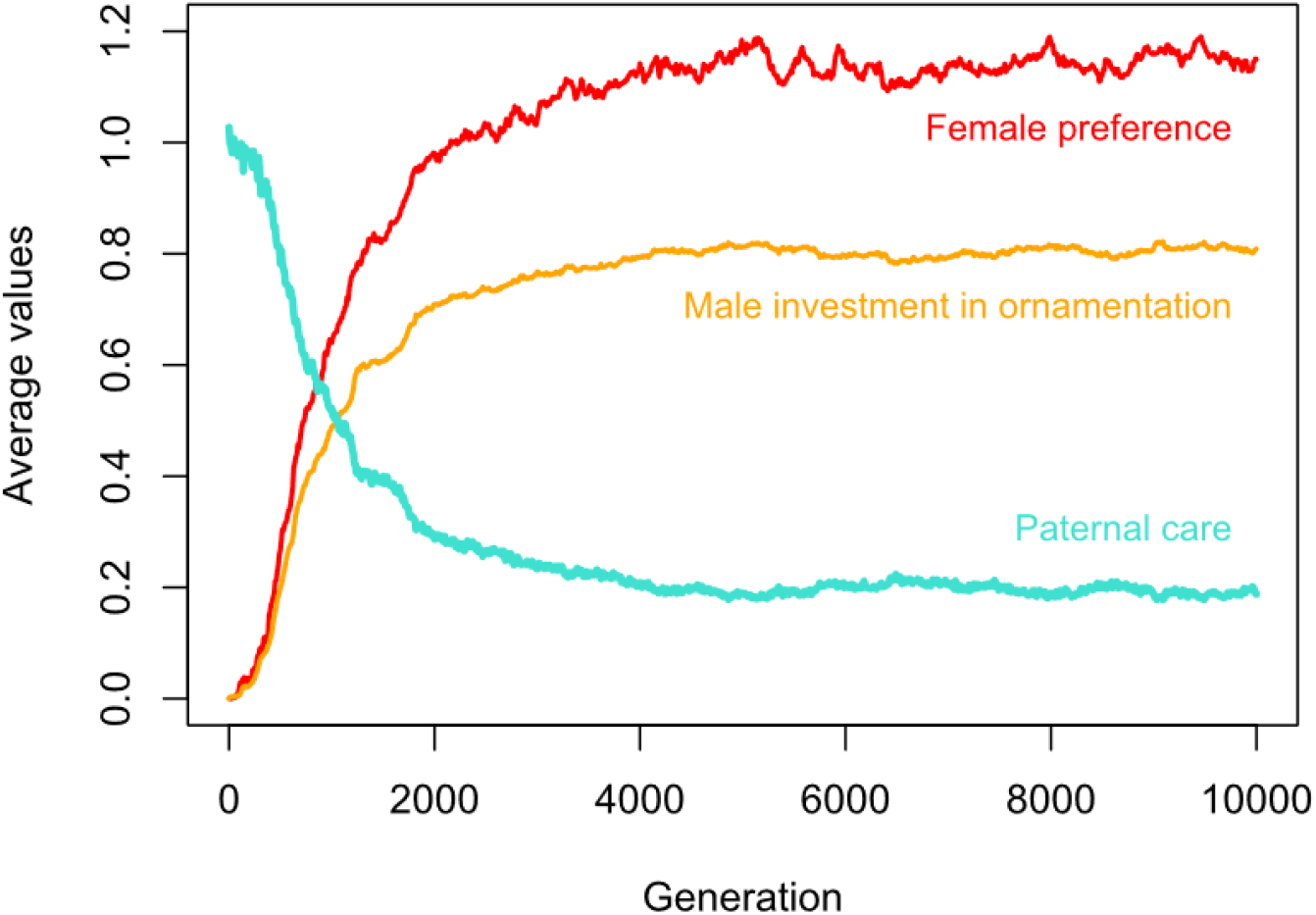
Evolution of female preferences, male ornamentation, and paternal care, shown for a representative simulation. Female preferences increase across generations (population mean female preference *P* shown in red); consequently, males evolve to invest more in ornamentation (population mean male investment in ornamentation *S* shown in orange), at the expense of paternal care (population mean paternal care shown in turquoise). Across generations, the co-evolution of female preferences and male ornamentation thus leads to the erosion of direct benefits (such as, in this case, paternal care). Parameter values are as in the baseline setting (long mating season and high resource variation), see methods.

### Sexually selected ornaments honestly signal direct benefits

Figure 3 demonstrates why females are selected to prefer more ornamented males, even though elaborate ornamentation reduces a male’s investment in paternal care. Since ornamentation is costly in terms of resources, only males with high resource levels produce large ornaments. These males also tend to provide more paternal care, that is, they provide a greater absolute amount of care, even if this may represent a smaller proportion of their resources: they invest a “smaller slice of a bigger pie”. Thus, both male resource level and paternal care level show a positive correlation with ornament size, throughout the simulation: male ornamentation remains an honest signal of paternal care (illustrated in Fig. 3A, for the same simulation that is shown in Fig. 2). Accordingly, females who preferentially mate with more ornamented males secure increased paternal care for their offspring, resulting in increased fecundity (Fig. 3B, Supplementary Fig. S1): even though offspring survival probabilities are overall declining, females consistently have better surviving offspring when mating with their chosen (ornamented) male rather than with a random male, throughout the course of the simulation. In other words, the ornament is an honest signal of paternal care, even though the ornament and paternal care trade off against each other.

**Figure 3:**
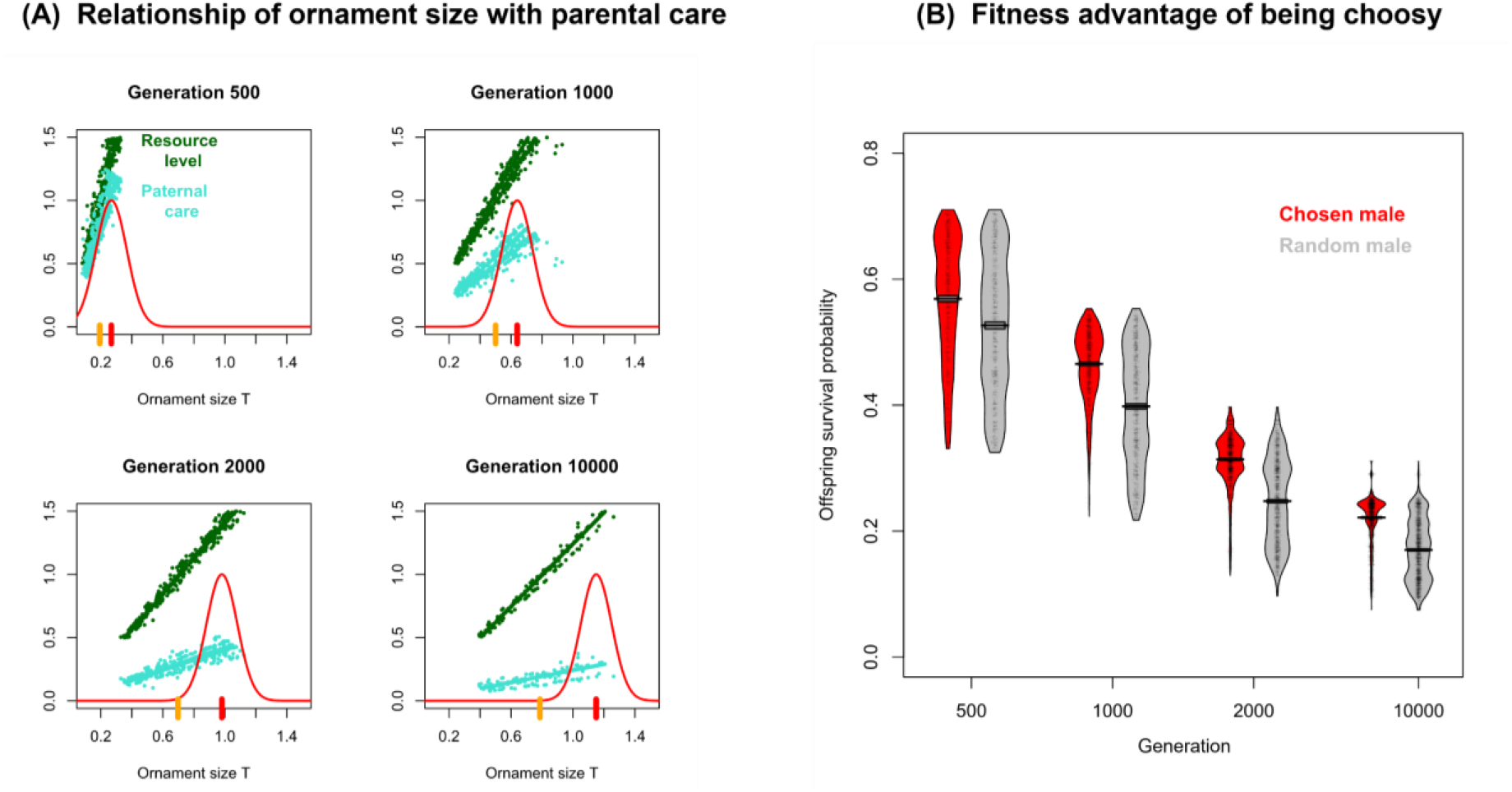
Detailed analysis of the simulation in Figure 2. **(A) Relationship of ornament size with resource level and paternal care**. Snapshots from four different generations show for all males in the population how their resource level (dark green) and the resources left for paternal care (turquoise) are related to their ornament size. At each time point, resource level and paternal care level (i.e., the direct benefits) are positively correlated with ornament size; hence, the ornament can serve as a reliable, honest signal of paternal quality. The red line indicates the probability of a female accepting a potential mate depending on his ornament size; coloured tick marks indicate the mean ornament size (orange) and female ornament size preference (red) in the population. As mean preference values are larger than mean ornament sizes, females exert positive selection on ornament size. Accordingly, both ornament sizes and preferred values increase over the generations. **(B) Fitness advantage of being choosy**. For the four time points in (A), the red violin plots show the survival probability of the offspring produced by the males that were chosen as mates by the females. For comparison, the grey violin plots show the survival probability of the offspring had they been the result of random mating. At all time points, mean offspring survival (horizontal bars) is higher under female choice than under random mating (confirming that, on average, females profit from being choosy). However, offspring survival declines substantially over the generations, due to the progressive erosion of paternal care.

### The evolutionary outcome depends on the length of the mating season and the extent of environmental resource variation

The outcome of the “baseline” scenario, as shown in Fig. 2, is consistent across replicate simulations (Fig. 4A): female preferences for larger ornaments (that indicate increased resource availability for paternal care) evolve; males evolve to invest in ornamentation at the expense of paternal care; and paternal care is thus progressively reduced. However, this dynamic is hindered if females have reduced opportunities to meet potential mates, such as due to short mating seasons (Fig. 4B): when females have time to encounter only two (rather than ten) males, females with high preferences are less likely to find a satisfactory mate. This limits the evolution of female preferences, leading to lower female preferences, lower investment in ornamentation, and higher levels of paternal care compared to the baseline scenario (Fig. 4B top). Still, ornamentation remains a reliable indicator of prospective paternal care, as larger ornaments are still associated with higher care levels (Fig. 4B bottom).

**Figure 4:**
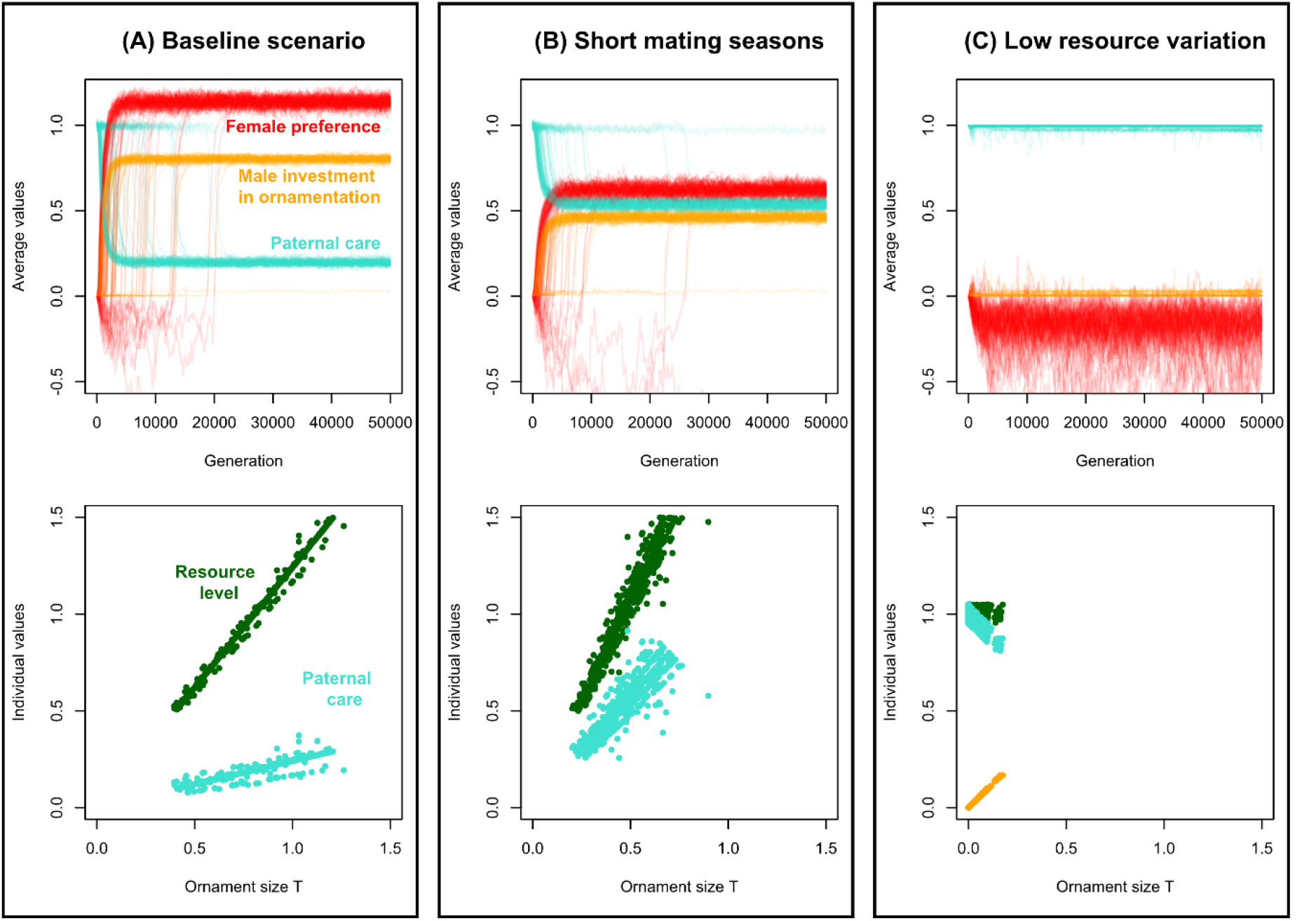
Dynamics of direct-benefits sexual selection in three scenarios. Top row: mean values of female preferences (red), male investment in ornamentation (orange) and the direct benefit (paternal care; turquoise) for 100 replicate simulations. Bottom row: the relationship of male ornament size with both the resources available to this male (dark green) and the paternal care provided by this male (turquoise), in a single representative replicate, at generation 10,000. Each point corresponds to a single individual. (**A) Baseline scenario**. Top: Female preferences increase, males evolve to invest in ornamentation at the expense of paternal care, and paternal care is strongly reduced. Bottom: Male ornamentation positively correlates with paternal care; it is thus an honest signal of mate quality. (**B) Short mating season**. Top: The increase in female preference and male investment in ornamentation is still present, albeit not as pronounced. Consequently, paternal care is maintained at a higher level. Bottom: male ornamentation positively correlates with prospective paternal care; it is thus an honest signal of mate quality. All parameter values are as in the baseline scenario, except for the maximal number of mate encounters, which was reduced from 10 to 2, to simulate a shorter mating season. (**C) Low resource variation**. Top: Female preferences evolve to be negative, i.e. females avoid ornamented males; males invest exclusively in paternal care. Paternal care is thus not eroded. Bottom: increased ornament size is associated with decreased prospective paternal care; higher mate quality is thus honestly signalled by the absence of ornaments. This reversal of the pattern seen in the first two scenarios (panels A and B) is due to variation in ornamentation being driven by variation in S (shown in orange) rather than variation in resource level. All parameter values are as in the baseline scenario, except that the range of resource variation has been reduced from 1 to 0.1 (see Methods).

Under low environmental resource variation (where resources are drawn from the interval [0.95,1.05], Fig. 4C), the evolution of female preferences for exaggerated ornamentation is fully suppressed: female preferences evolve to be negative (i.e. females avoid, rather than seek out, ornamented males), and males maintain full investment in paternal care, at the expense of any investment in ornamentation (Fig. 4C top). Variation in male ornamentation is driven both by resource variation (under which ornamentation and paternal care are positively correlated) and by variation in the male investment strategy *S* (under which ornamentation and paternal care are negatively correlated). Thus, with decreasing resource variation, variation in ornamentation increasingly reflects variation in *S* – the signal not only loses its reliability but, under low resource variation, becomes associated with *decreased* levels of paternal care (Fig. 4C bottom). Females thus benefit from avoiding ornamented males, as high paternal care levels are here indicated by the absence of ornamentation, leading to the evolution of negative preferences. We show here two scenarios at the extremes of the resource variation spectrum: very broad (Fig. 4A) and very narrow (Fig. 4C) variation. For simulations of intermediate levels of resource variation, we found that replicates differed in whether they evolved towards a “sexual-selection-scenario” (high ornamentation, low paternal care) or towards a “paternal-care-scenario” (no ornamentation, full paternal care; see Supplementary Fig. S2).

### Direct-benefits sexual selection can drive population extinction

For all results shown so far, we assumed that the size of the population is fixed. In nature, this is an unrealistic assumption. We thus test whether our results are replicable when population sizes are not fixed, but can fluctuate depending on offspring survival (which, in turn, depends on the degree of parental care received, thereby linking evolutionary and ecological dynamics). The evolutionary dynamics under fluctuating population sizes is the same as under fixed population sizes (Fig. 5A): females preferring ornamented males have, on average, more offspring, and thus, over time, male ornamentation and female preferences for indicators of paternal care increase, whereas paternal care and offspring survival decrease. However, once average male investment in paternal care falls below a certain threshold value, the population can no longer sustain itself and goes extinct. In other words, sexual selection drives population extinction. This pattern is again highly repeatable across replicates (98 out of 100 replicates went extinct within 50K generations). In scenarios where the evolution of extreme female preferences and male ornamentation is impeded, such as in environments with short mating seasons (Fig. 5B) or little resource variation (Fig. 5C), extinction is reduced as well (20 out of 100 replicates show extinction under short mating seasons, 0 out of 100 replicates show extinction under little resource variation within 50K generations).

**Figure 5:**
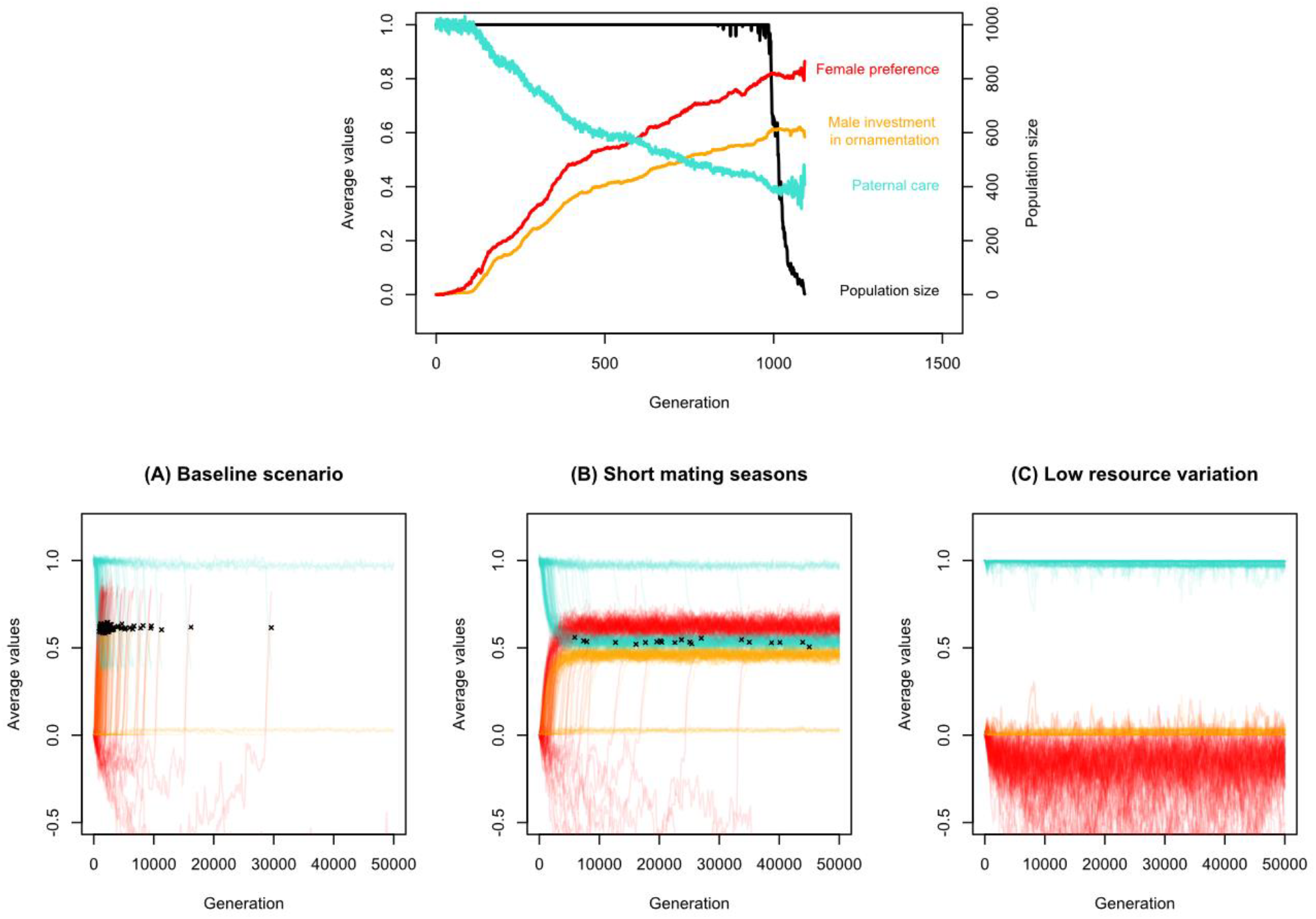
Population dynamics under direct-benefits sexual selection. The three scenarios in Fig. 4 are considered again, but now with a variable population size. Top row: extinction dynamics in a representative simulation of the baseline scenario. Bottom row: Extinction dynamics under different scenarios, showing mean values of female preferences (red), male investment in ornamentation (orange) and the direct benefit (paternal care; turquoise) for 100 replicate simulations. Each extinction event is marked by a black cross at the end of the line that shows the mean male investment in ornamentation. **(A) Baseline scenario**. Female preferences and male investment in ornamentation increase, and paternal care decreases. As offspring survival depends on paternal care, this eventually leads to population extinction in almost all replicates, as not enough offspring survive to replace the parental generation. **(B) Short mating season**. When the number of mating opportunities is low, females evolve preferences for ornamented males, but these are lower than in the baseline scenario. Consequently, males evolve to invest in both ornamentation and paternal care, and extinction occurs less frequently than in the baseline scenario. **(C) Low resource variation**. When resource variation is low, females evolve preferences for unornamented males and males evolve to invest resources in paternal care rather than ornamentation. Extinction is thus prevented.

### Reduced importance of male care destabilises sexual selection

The model versions underlying Fig. 2-5 assume that paternal care is essential for offspring survival; that is, without paternal care, the offspring cannot survive. This is the case for systems with male-only care or for systems with biparental care where the care contributions of one parent alone are insufficient for offspring survival. Under these conditions, we see that the dynamics of sexual selection reach an equilibrium (as long as the population does not go extinct first). Similar results are obtained when offspring survival without paternal care is low (rather than zero). For a survival probability of 0.2 in the absence of male care, Figure 6B shows that female preferences and male ornamentation converge to even higher levels, reducing parental care to even lower levels than in our baseline scenario (Fig. 4A, Fig. 6A). However, the evolutionary dynamics changes fundamentally if offspring can survive well without paternal care. For a survival probability of 0.5 in the absence of male care, this is exemplified by the pattern exhibited by the 100 replicate simulations in Figure 6C, which looks more complicated than the pattern in Fig. 6A-B. The time plot of a representative simulation in Figure 6D indicates a systematic sequence of events: Initially, female preferences and male ornamentation increase at the expense of paternal care. However, after a while, this “high-ornamentation-low-care” state collapses, and males return to investing predominantly in paternal care rather than ornamentation. After some time, this newly achieved “low-ornamentation-high-care” state also collapses, and the system returns to the state with low paternal care. Figure 6E shows that this pattern of repeated switching between a quasi-equilibrium with high ornamentation and low parental care and a second quasi-equilibrium with low ornamentation and high parental care occurred in a stochastic but systematic manner. In other words, the system does not converge to equilibrium when offspring survival is high; instead, individual trajectories are highly labile and repeatedly switch between two states, rapidly gaining or losing ornamentation between prolonged periods of relative stasis.

**Figure 6:**
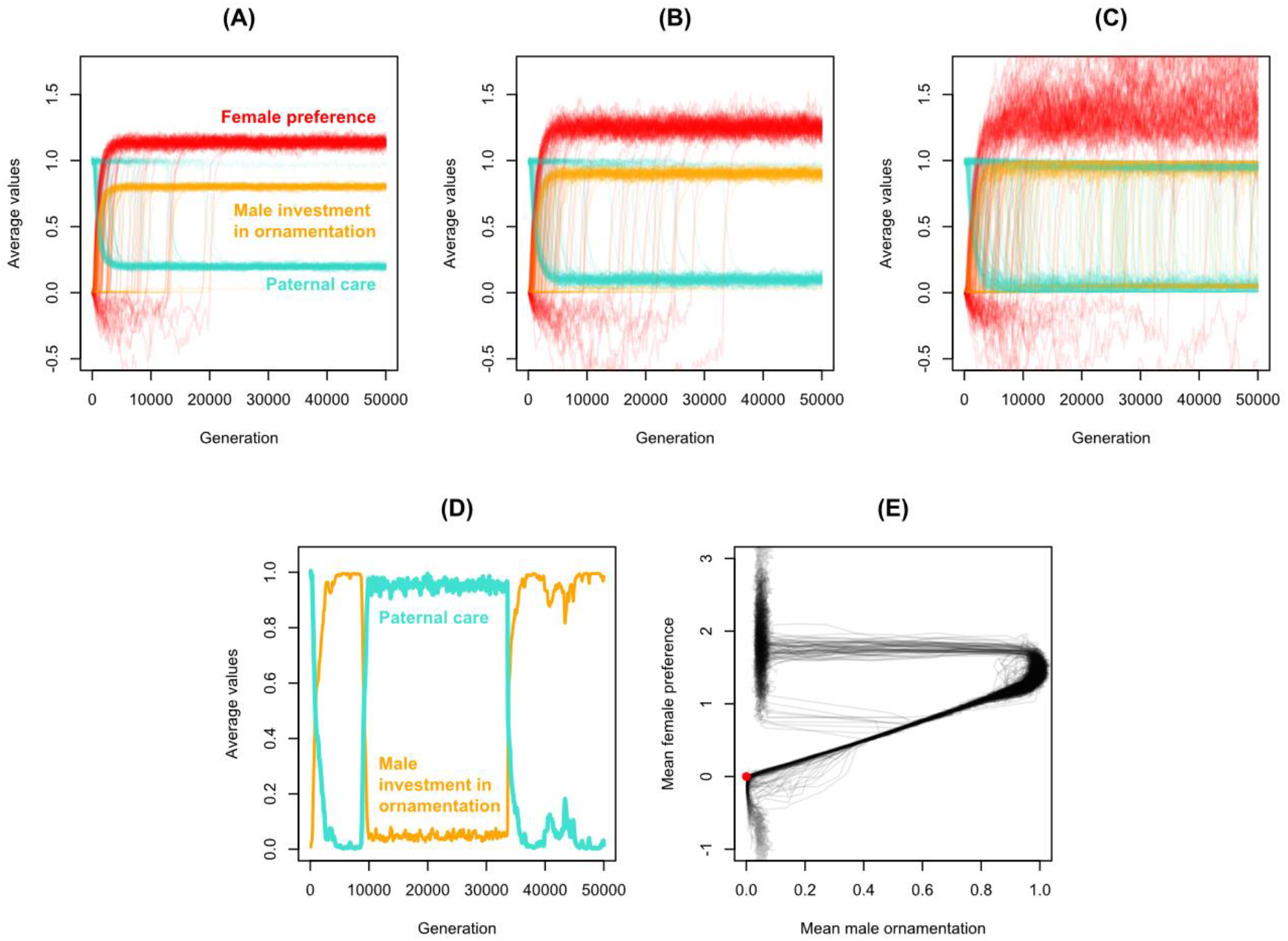
Dynamics of direct-benefits sexual selection under different parental care scenarios. Mean values of female preferences (red), male investment in ornamentation (orange) and the direct benefit (paternal care; turquoise) are shown for 100 replicate simulations (A-C). **(A) No offspring survival without male care** (as in Fig. 4A). Female preferences and male investment in ornamentation increase, and paternal care decreases. An equilibrium value for these traits is reached. **(B) Low offspring survival without male care**. Similar to the baseline scenario, equilibrium values for female preferences, male investment in ornamentation, and male care are reached. Mean female preferences and male investment in ornamentation are higher than in the baseline scenario; correspondingly, mean male care levels are lower. **(C) High offspring survival without male care**. Similar to the baseline scenario, female preferences and male investment in ornamentation initially increase, at the expense of paternal care. However, sexual ornamentation cannot be maintained but is repeatedly lost and regained. **(D) A single representative simulation** under high offspring survival without male care. Ornamentation repeatedly evolves, is lost, and regained, as the system rapidly and abruptly switches between two states: one in which males invest almost all resources in ornamentation, and one in which males invest almost all resources in paternal care. **(E) Phase plot depicting mean female preference plotted against mean male ornamentation**, evolving for 50K generations under high offspring survival without male care. The red dot indicates the mean values at the start of the simulation; the mean values for all 100 simulations are shown. After an initial increase in both female preferences and male ornamentation, the system switches between two quasi-equilibria: one with high ornamentation and low parental care (not shown here) and one with low ornamentation and high parental care. All parameter settings are as in the baseline scenario in Fig. 4A; the population sizes are fixed.

## Discussion

In our model, male ornamentation provides a reliable signal of prospective paternal care (i.e. of the direct benefit a female may receive), and females consequently readily evolve preferences for more ornamented males. However, these preferences in turn select for more ornamented males – and increased investment in ornamentation trades off against investment in paternal care: sexual selection fuelled by the acquisition of direct benefits thus erodes direct benefits. These results, though consistent across replicates, depend on sufficient opportunities for female choice, as well as on sufficient variation in male quality. If opportunities for female choice are reduced, female preferences and male ornamentation evolve to be less exaggerated. If males do not vary sufficiently in quality (i.e., in the resources they provide), females do not evolve preferences for ornamented males; consequently, males evolve to invest in paternal care rather than ornamentation. In summary, given sufficient opportunity for mate choice and variation in mate quality, the dynamics of direct-benefits sexual selection, though readily established, erode direct benefits.

In our model, we consider a scenario where the cost of ornamentation is reduced fecundity: males that invest more in ornamentation have fewer resources left for paternal care and thus produce fewer surviving offspring per mating. This assumption is crucial to our results, and it is arguably the most important point of departure from most previous models of direct-benefits sexual selection. Some earlier models assume that females can directly assess the direct benefits a prospective mate will provide, thus circumventing the need for (potentially costly) signals^25–29^. Other models do assume that males signal their direct benefits to potential mates using costly ornaments, but assume that these costs are paid by the males in terms of reduced viability^35^. Both types of model (no signalling costs and viability costs of signalling) exhibit similar dynamics: females evolve preferences for males who provide more direct benefits; as females obtain more direct benefits, female fecundity increases. Specifically, Kirkpatrick shows that at equilibrium, female fecundity is maximised^25^. Our assumption that the costs of ornamentation are paid in terms of reduced fecundity is fundamentally different, as a fecundity cost is by necessity paid by both parents: females who mate with more ornamented males thus not only profit from this male’s direct benefits but also share in the cost of ornamentation. This is at the heart of the findings from our model: if both sexes carry the cost of ornamentation, then this cost compromises the direct benefit a female may gain from mating with ornamented males, and female fecundity declines even as her preference for males that provide direct benefits increases. Our results are in line with the quantitative genetics model in Price et al. (1993)^34^, who also considered a fecundity cost of ornamentation, and similarly showed that female fecundity may decrease under direct-benefits sexual selection. To the best of our knowledge, this is the only other dynamic evolutionary model considering a cost carried by both parents (but see the verbal model of Fitzpatrick 1995^44^ and the game-theoretical analysis of Kelly & Alonzo 2009^45^ for conceptually similar approaches).

When the availability of direct benefits is signalled through sexual ornamentation, what ensures that this ornamentation is an honest signal? Previous direct-benefits models mostly link ornamentation and direct benefit by either presupposing a (positive) correlation between both or modelling the ornament as condition-dependent^34–36^. We take an alternative route (following Kokko 1998^46^ and Kelly & Alonzo 2009^45^), by assuming that ornamentation and benefit are linked because both of them represent a portion of a shared resource pool: for a given level of proportional investment, males with more resources will produce both a larger ornament and provide more paternal care. As long as males do not significantly differ in the proportion invested in ornamentation, females can then use a male’s ornamentation to obtain a reliable estimate of a male’s resource level, and consequently also of the male’s prospective paternal care level. Thus, even though ornamentation and paternal care trade off against each other, the level of ornamentation can be positively correlated with the level of care: males with more resources have both bigger ornaments and provide more care.

However, ornamentation is not reliably associated with increased care under some circumstances. This occurs under low resource level variation: here, the variation in male investment strategy is relatively high compared to resource variation, and thus larger ornaments are associated with lower paternal care (rather than higher resources and therefore higher care). This illustrates a general principle in life history theory: the importance of distinguishing between acquisition and allocation^47^. In essence, this principle states that two quantities that draw on the same resource pool can be positively or negatively correlated, depending on whether variation in these quantities is predominantly driven by variation in resource acquisition or by variation in resource allocation. In our model, ornamentation and paternal care trade off against each other, as they draw on the same resource pool of an individual. If males allocate resources in the same way but differ in how much they acquire, then the system is acquisition-driven: those with more resources produce larger ornaments and provide more care, and the two traits are therefore positively correlated. Conversely, if males acquire similar amounts of resources but differ in how they allocate them, then the system is allocation-driven: males investing more in ornamentation invest less in care, and the two traits are therefore negatively correlated. If we now reconsider the question of signal honesty in the light of acquisition vs. allocation, we can see that, despite of the trade-off between the signal and the paternal care level to be signalled, the signal can still be honest as long as the system is acquisition-driven (as is the case in our baseline scenario); however, if the system is allocation-driven (as is the case when environmental resource variation is too low, leading to reduced variation in resource acquisition), higher signal magnitudes are associated with lower paternal care.

There is a large and strongly divided body of empirical literature on whether, in nature, parental care and sexual traits are generally positively or negatively correlated. On the one hand, the literature on the “good parent process”^41^ – where males’ ornaments signal their prospective levels of care – provides examples of positive correlations between ornamentation and offspring care (e.g. ^48–52^). On the other hand, many taxa exhibit negative correlations between ornamentation and offspring care (e.g.^53–55^, reviewed in ^56^ and for bird taxa in ^57^). Several explanations have been put forward for such negative correlations. First, as in our model, they may reflect fundamental allocation trade-offs, since energy, resources, or time invested in mate attraction often cannot also be invested in offspring care (reviewed in ^56^). Second, individuals may plastically adjust their care level depending on their own or their mate’s attractiveness, as predicted by the differential allocation hypothesis (females mated to attractive males may work harder, thus decreasing the necessity of paternal care^43^): males that are less attractive may increase their offspring care level to invest more in the current brood, since they likely have fewer future breeding opportunities (e.g.^40^). Overall, empirical evidence for both positive and negative correlations between ornamentation and offspring care is abundant; meta-analyses and literature reviews assessing their relative prevalence suggest no clear overall pattern but considerable variation across studies and species; and conflicting evidence is not only reflective of taxonomic variation: different studies on the same species repeatedly find opposing patterns (reviewed in^13,56–58^). We suggest that our model, together with an understanding of acquisition-driven vs. allocation-driven variation, may shed some light on this. Specifically, within a generation, acquisition-driven positive correlations between ornamentation and parental care allow ornamentation to function as an honest signal. This drives greater investment in ornamentation, and thus less investment in care. Across evolutionary time, therefore, ornamentation and parental care are negatively correlated due to variation in allocation strategy – males at later timepoints invest more in ornamentation than males at earlier timepoints. Thus, empirical studies of honest signals of paternal care may find (even within the same study system) positive correlations when assessing correlations within generations or populations, but negative correlations when assessing correlations across populations, species, or longer evolutionary timescales. Similar arguments have been made in the context of other life-history trade-offs (such as the trade-off between current and future reproduction)^59^. In line with this, future meta-analyses may be able to investigate to what extent conflicting patterns in the empirical literature on parental care and sexual traits can indeed be explained by the scale at which their correlation is assessed.

When population sizes are not kept fixed, we find that many simulation replicates “evolve towards extinction”: as selection drives investment in ornamentation at the expense of paternal care, offspring survival declines, and the population goes extinct. Previous research (reviewed in^60–62^) has explored the effects of sexual selection on population viability, with findings ranging from sexual selection facilitating adaptation and improving population viability (e.g.^63,64^), to sexual selection acting at cross-purposes to natural selection and hindering adaptation (e.g.^7,8^), to sexual selection driving extinction (e.g.^9,65,66^). Our results fall into the latter category. However, in previous examples of such “evolutionary suicide”, extinction is the consequence of a maladaptive runaway process, often involving arbitrary preferences. In contrast, extinction in our model occurs at evolutionary equilibrium and results from female preferences for honest indicators of male fecundity. The possibility that direct-benefits sexual selection might lead to fertility decline and reduced population viability was already discussed in earlier work^34,44^. Yet, to our knowledge, this is the first study to explicitly demonstrate population extinction driven by direct-benefits sexual selection.

Although we, for illustrative purposes, refer to paternal care as the direct benefit that males provide to females, our results generalise to other types of direct benefits (e.g. access to better territories, nuptial gifts that improve female condition, or defence against predators, which also increase offspring survival). Nevertheless, some assumptions in our model may not apply to all types of direct benefits. For instance, we assume that there is an inherent trade-off between investment in the direct benefit and investment in ornamentation; this is plausible if the direct benefit is parental care (as ornamentation may reduce the male’s ability to forage food for the offspring, or because males cannot simultaneously feed the offspring and perform sexual displays), but it may be less plausible if the direct benefit is access to a good territory (as the same ornamentation or sexual display that attracts females may also aid the male in defending this territory against rival males; for more counterexamples see^56^). Similarly, we assume for simplicity that all offspring of a male benefit equally from the male’s investment, irrespective of the number of offspring fathered by the male (Kirkpatrick’s 1985^25^ “unlimited male reproductive potential” scenario). This assumption is plausible if the direct benefit is a reduced risk of disease transmission, but less so if the male must feed all his offspring. We also made two other assumptions worth noting. First, in our model, male “quality”, i.e. resource availability, is entirely environmentally determined. This assumption was made deliberately, as we aimed to investigate the dynamics of direct-benefits sexual selection, and thus attempted to exclude other processes, such as good-genes sexual selection. If males differ genetically in their ability to provide direct benefits, good-genes and direct-benefits sexual selection become closely intertwined, making it virtually impossible to disentangle their relative contributions. However, the investigation of such an intertwined model led to conclusions that are very similar to those of the present study^67^.

Second, in our model, we consider several implementations of male-only or biparental care (Fig. 6). When male care is not sufficiently important for offspring survival (as may the case in e.g. a predominantly female care system), we find that the evolutionary dynamics is destabilised, and the system oscillated between two quasi-equilibria (characterized by high ornamentation/low care and low ornamentation/high care respectively). Similar patterns have also been described for other forms of sexual selection^10,66^, and are reflected in empirical data: sexual ornaments are often phylogenetically labile and show high rates of turnover^68^. However, in our implementations of parental care, we abstract away two aspects of female offspring care (evolution): female investment in offspring care may co-evolve with female preferences and male care^39,69^, and females may plastically adjust their care levels based on characteristics of their mate, either to (partially) compensate for reduced male care or to increase investment in the more “promising” offspring produced with a high-quality male^43,70–72^. Future models should explore how the dynamics of direct-benefits sexual selection are impacted by plastic or evolutionary changes in female care in response to the evolution of male care.

Using individual-based evolutionary simulations, we show that direct-benefits sexual selection is far more complex than often assumed. Standard analytical models are based on simplifying assumptions such as a monomorphic population. Individual-based simulations provide a flexible and easily extendable way to model complex scenarios where individual interactions can be implemented in a more natural way^20^. Using such a model, we showed how the interplay of resource acquisition and allocation leads to a scenario where the selection on female preferences to maximise offspring survival results in decreased offspring survival. Thus, direct-benefits sexual selection leads to population extinction through female preferences for honest indicators of male fecundity. These results lead us to conclude that the theory of direct-benefits sexual selection deserves more attention than it currently receives and that, with evolutionary simulations that consider more realistic assumptions, even apparently straightforward scenarios can lead to unexpected and counterintuitive results.

## Methods

### Model structure

We model a population of *N* = 1000 individuals with discrete, non-overlapping generations, evolving through rounds of mating and reproduction. Each male is randomly assigned a resource level *R* at birth. Based on a heritable strategy *S*, each male decides what fraction of *R* to devote to an ornamental trait T and what fraction to retain for the direct benefit *B*. Each female has a heritable mating preference *P*, which determines the probability of accepting a male with trait *T* for mating. A mated pair produces one offspring; the survival probability of the offspring is positively related to the benefit *B* provided by the father. Females and males can mate multiple times; the mating process continues until *N* surviving have been produced. Offspring are assigned a sex at random, and they inherit the alleles underlying the heritable strategies *S* and *P* from their parents in a Mendelian fashion, subject to rare mutations. Once *N* surviving offspring have been produced, the parental population dies and is replaced by the offspring population. This process is repeated for 50,000 generations. Below, we describe a model variant in which the population size is not constant but variable; in this case, the population can decline (potentially to extinction) if offspring survival is very low.

### Resource assignment, male investment strategy, and offspring survival

At birth, each male is endowed with a resource level *R. R* is randomly drawn from a uniform distribution that is centred on one and has a range of 2*D*. Hence, *R*_min_ = 1− *D* and *R*_max_ = 1+ *D* By default, we chose the value *D* = 0.5. However, *D* was reduced to *D* = 0.05 in the “low resource variation” scenario.

The individuals in our population are haploid (see Supplementary Figure S3 for the outcome of a model variant with diploid individuals) and characterised by the alleles at two gene loci: the *S*-locus (expressed only in males) and the *P*-locus (expressed only in females). The alleles at the *S*-locus correspond to real numbers from the unit interval [0,1] and determine the fraction of resources invested in ornamentation. Hence, a male *i* endowed with resource level *R*_*i*_ and harbouring allele *S*_*i*_ develops an ornament of size *T*_*i*_ = *S*_*i*_ · *R*_*i*_. The male’s investment in his ornament trades off against the direct benefits *B*_*i*_ provided by the male (e.g. the level of paternal care), such that *B*_*i*_ = (1 − *S*_*i*_) · *R*_*i*_. Accordingly, both ornamentation and direct benefits are proportional to the resource level (a more general allocation strategy, leading to similar results, was investigated in Mosna 2017, MSc thesis, Univ. of Groningen). The survival probability of the offspring fathered by male *i* is given by 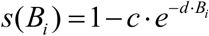 and therefore increases with *B*_*i*_ (the degree of paternal care received). The parameters *c* and *d* are scaling factors determining the shape of the relationship between investment in offspring care and offspring survival. By default, the simulations reported in the main text are based on the setting *c* = *d* = 1. The parameter *c* determines how strongly offspring survival depends on male care. Specifically, 1 *c* gives the probability of offspring survival in the absence of male care. We chose *c* = 0.8 and *c* = 0.5 for the scenarios of biparental care considered in Fig. 6.

### Mate choice and reproduction

The allele at the *P*-locus determines the females’ mating preference. In line with Lande’s (1981)^73^ “absolute preference” model, we assume that a female *j* with allele *P*_*j*_ at the *P*-locus accepts a male with trait *T*_*i*_ for mating with probability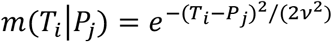. This mating function is bell-shaped with a peak at *T*_*i*_ = *P*_*j*_, which implies that males whose trait matches the female preference have the highest mating probability. The parameter ν determines the width of the mating function and, hence, the female’s tolerance for mating with males with a trait deviating from her preference. As a rule of thumb, males have a high probability of being accepted for mating if their trait value deviates less than ±ν from the female’s preference. In our simulations, we used the value ν = 0.1.

We modelled the mating process as follows: In a stepwise fashion, females are chosen at random from the population. Each chosen female encounters one or several males until she accepts a male for mating. One by one, males are chosen at random; if *P*_*j*_ is the female’s preference and *T*_*i*_ is the male’s trait, the probability of mating is given by *m*(*T*_*i*_|*P*_*j*_). If the female rejects the male, she will encounter another male (again selected at random from the population). This is repeated until the female is either mated with a preferred male or until the female has encountered *M* males, in which case she mates with the last (random) male. For example, when *M* = 10, if the female rejects the first nine males, she mates with the tenth, regardless of his ornament. This assumption mimics the decline in female choosiness as the breeding season progresses, which has generally been interpreted as the female’s “lowering of standards” to avoid remaining unmated ^74,75^. By default, we chose the value *M* = 10. However, *M* was reduced to *M* = 2 in the “short mating season” scenario. Once a female has accepted to mate with a male, the mated pair produces one offspring, which survives with probability *s*(*B*_*i*_) where *B*_*i*_ is the benefit level provided by the offspring’s father. In the next step, a new female is chosen at random and the same procedure is followed until *N* surviving offspring have been produced.

### Inheritance

Each surviving offspring is assigned a sex (female or male) at random. Male offspring are provided with a resource level *R* as described above. Moreover, each offspring inherits alleles at the *S*- and *P*-locus from its parents. At each locus, it is decided independently whether the maternal or the paternal allele is transmitted to the offspring. Each of the transmitted alleles mutates with probability *μ* = 0.01. If a mutation occurs, the parental allele *A*_*k*_ is changed to *A*_*k*_ + *δ*, where the mutational step size *δ* is drawn from a normal distribution with mean zero and standard deviation 0.05. As the alleles at the *S*-locus correspond to a fraction (and are therefore limited to the unit interval [0,1]), a new mutational step size is drawn whenever a mutated allele *A*_*k*_ + *δ* at the *S*-locus is either smaller than zero or larger than one. In all simulations presented, the alleles at the *S*- and *P*-locus were initialised at the value zero for all individuals in the population.

In Supplementary Figure S3, we consider a diploid variant of the model. If we make the standard assumption that the *S*- or *P*-strategy of an individual corresponds to the average of the two alleles at the *S*- or *P*-locus, the simulation results are virtually identical to those of the haploid model considered in the main text.

### Variable population size

To explore the impact of sexual selection on population persistence, we developed a model version where population size is not fixed. To do so, we implemented the following changes to the model described above. Instead of producing a single offspring with various males, each female mates only once. The mating process is as explained above: a female assesses up to *M* males, and if she rejects the first *M* − 1 males, she has to mate with the *M* ^th^ male. A mated pair produces *n* offspring, where the litter or clutch size *n* is drawn from a Poisson distribution with parameter 5. Hence, a mated pair produces 5 offspring on average. However, not all of these offspring survive: the survival probability per offspring is again given by 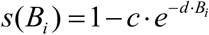, where *B*_*i*_ is the direct benefit provided by the father. Moreover, the previously fixed population size *N* = 1000 is now viewed as the carrying capacity of the population: if more than *N* surviving offspring are produced as described above, offspring are removed at random until the number of surviving offspring no longer exceeds the carrying capacity *N*.

## Data and code availability

The C++ code for the simulations, the simulated data used to generate figures in this study, and the R code for data analysis can be found at https://github.com/Jana17/DirectBenefitsSexualSelection_2025.

## Acknowledgements

F.J.W. and J.M.R. acknowledge funding from the European Research Council (ERC Advanced Grant No. 789240). J.M.R. was also supported by a GELIFES scholarship from the University of Groningen. We thank Flavia Di Santo for the drawings used in Figure 1.

