## Supplementary Material for "Sexual selection driven by direct benefits leads to the erosion of direct benefits"

**Contents**

**Figure S1:** Offspring survival in the case of a choosy mother and a randomly mating mother.

**Figure S2:** The dynamics of direct-benefits sexual selection under different degrees of resource variation.

**Figure S3:** Comparison of model implementations based on haploid and diploid inheritance.

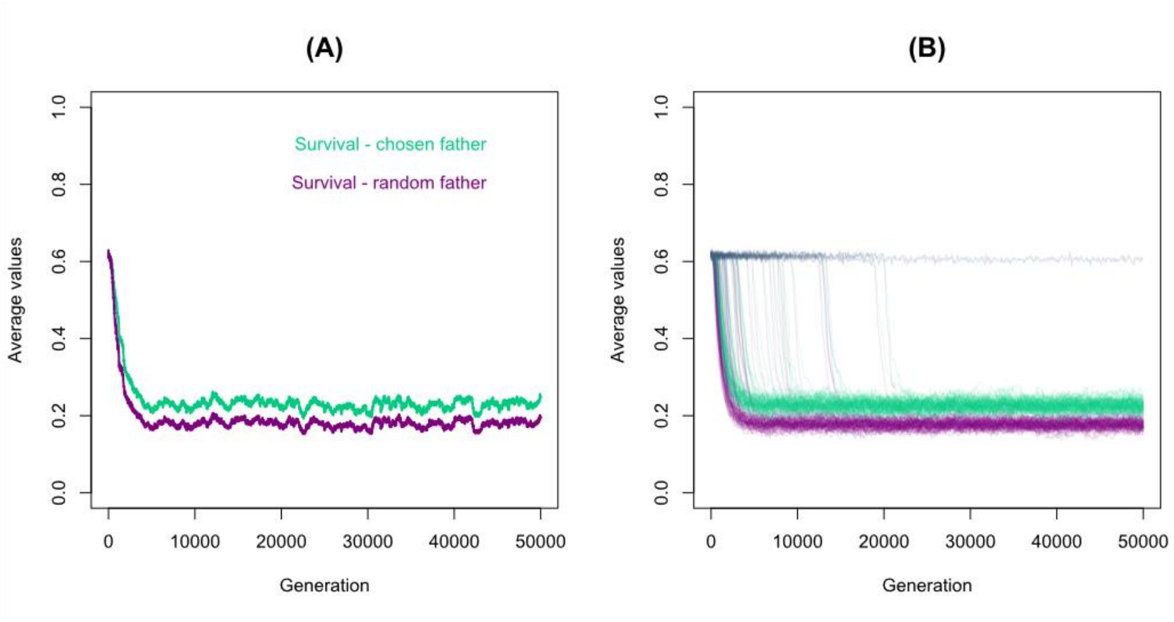


**Figure S1: Offspring survival in the case of a choosy mother and a randomly mating mother**, shown for **(A)** a representative simulation and **(B)** for 100 replicate simulations. Throughout the simulation, the offspring produced by females with their chosen mate have higher survival probabilities (green) than the offspring which the females would have produced if they had mated at random (purple): in other words, females benefit from being choosy. Nevertheless, average offspring survival declines over the course of the simulation. All parameter values are as in the baseline setting.

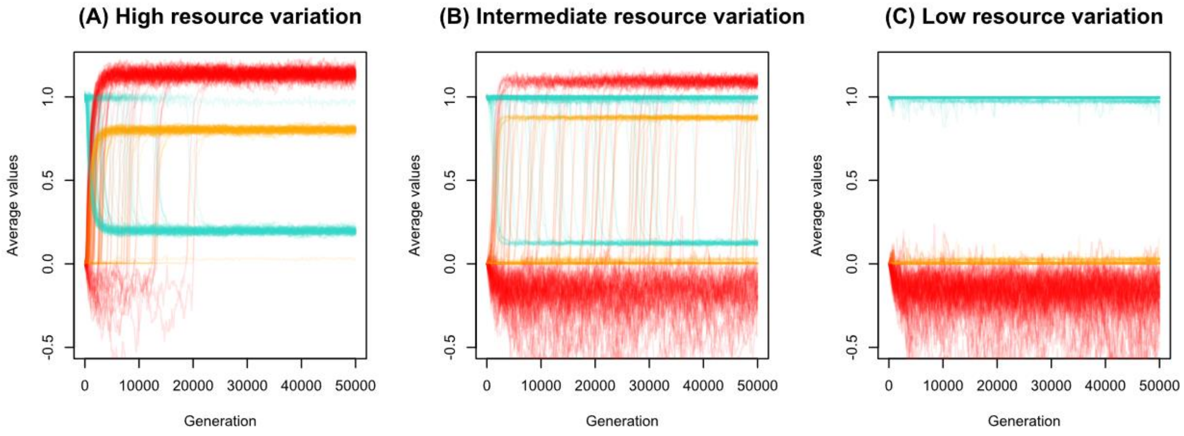


**Figure S2: Dynamics of direct-benefits sexual selection under different degrees of resource variation. (A)** When resource variation is high, as in the baseline scenario, male investment in ornamentation (shown in orange) and female preferences (shown in red) evolve to be high, at the expense of paternal care (shown in turquoise). **(C)** When resource variation is low, females prefer unornamented males, and males invest into paternal care and instead of ornamentation. **(B)** When resource variation is at an intermediate level, both outcomes can be observed: out of 100 replicate simulations, 52 evolve towards a "sexual-selection-scenario" (high ornamentation, low paternal care), whereas 48 evolve towards a "paternal-care-scenario" (no ornamentation, full paternal care). All parameter values except for the range of resource variation are as in the baseline setting; Resources are drawn from the interval [0.5,1.5] in **(A)**, from the interval [0.75, 1.25] in **(B)** and from the interval [0.95,1.05] in **(C)**.

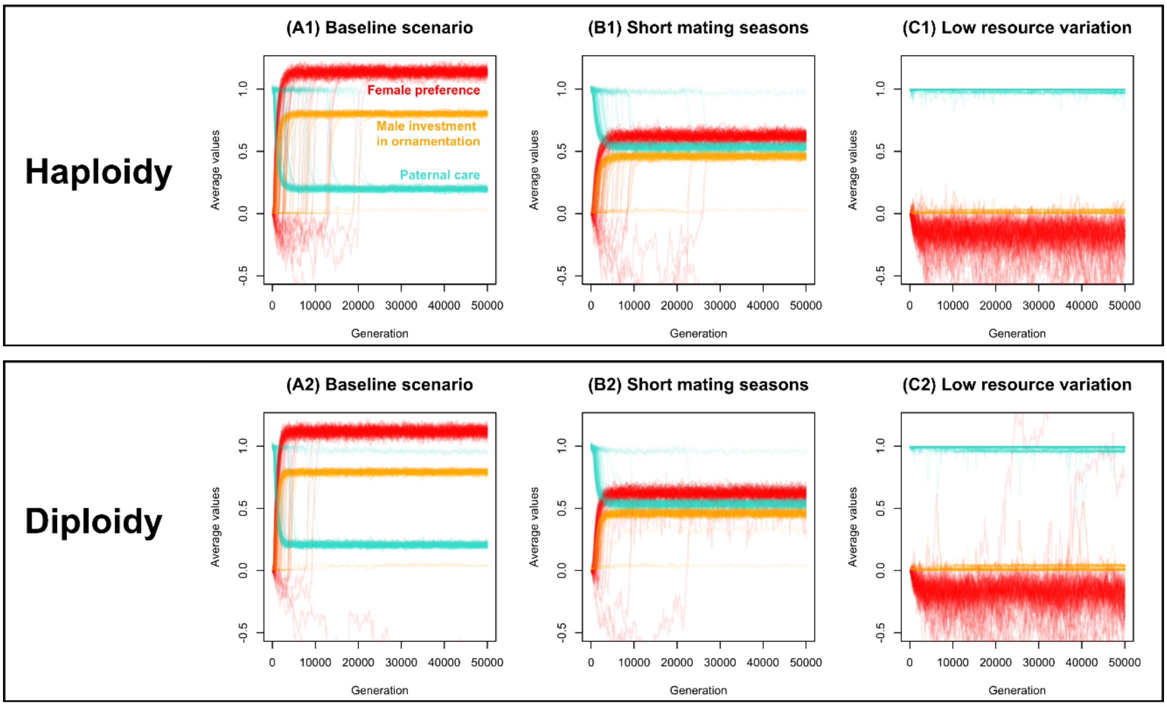


**Figure S3: Comparison of model implementations based on haploid and diploid inheritance.** The dynamics of direct-benefits sexual selection do not vary significantly between model implementations with haploid genetics **(A1, B1, C1)** and diploid genetics with additive gene effects **(A2, B2, C2)**. This holds true for all scenarios considered: the baseline scenario **(A)**, the scenario with short mating seasons **(B)**, and the scenario with low resource variation **(C)**. All other parameters are as in the baseline setting. Panels A1-C1 correspond to the panels shown in the top row in Fig. 4.
